# Cohesin interacts with a panoply of splicing factors required for cell cycle progression and genomic organization

**DOI:** 10.1101/325209

**Authors:** Jung-Sik Kim, Xiaoyuan He, Jie Liu, Zhijun Duan, Taeyeon Kim, Julia Gerard, Brian Kim, William S. Lane, William S. Noble, Bogdan Budnik, Todd Waldman

## Abstract

The cohesin complex regulates sister chromatid cohesion, chromosome organization, gene expression, and DNA repair. Here we report that endogenous human cohesin interacts with a panoply of splicing factors and RNA binding proteins, including diverse components of the U4/U6.U5 tri-snRNP complex and several splicing factors that are commonly mutated in cancer. The interactions are enhanced during mitosis, and the interacting splicing factors and RNA binding proteins follow the cohesin cycle and prophase pathway of regulated interactions with chromatin. Depletion of cohesin-interacting splicing factors results in stereotyped cell cycle arrests and alterations in genomic organization. These data support the hypothesis that splicing factors and RNA binding proteins control cell cycle progression and genomic organization via regulated interactions with cohesin and chromatin.

**One Sentence Summary:** Endogenous tagging reveals that cohesin interacts with diverse chromatin-bound splicing factors that regulate cell cycle progression and genomic organization in human cells.

## Main Text

Cohesin is a ubiquitously expressed multi-protein complex best known for its involvement in sister chromatid cohesion, but which also plays important roles in chromosome organization, gene expression, and DNA repair (1,2,3,4,5). Somatic mutations of cohesin are common in diverse cancers, and inherited mutations result in developmental disorders known as cohesinopathies (6,7). In vertebrate cells, cohesin is a ring-like structure encircling chromatin composed of four core subunits: SMC1A, SMC3, RAD21, and either STAG1 or STAG2. Several additional subunits serve to regulate the core complex, including NIPBL, MAU2, WAPL, PDS5A, PDS5B, and Sororin. The cohesin complex is highly conserved in both prokaryotic and eukaryotic unicellular organisms, as well as in metazoans.

To obtain a comprehensive picture of human cohesin protein-protein interactions, gene editing was used to add a dual FLAG-Streptavidin Binding Peptide (SBP) epitope tag to an endogenous allele of each of the genes encoding the 11 known components of cohesin in HCT116 cells, a human cell line with wild-type cohesin genes and intact sister chromatid cohesion (Table S1, Fig. S1-S11) (8). AAV-based gene editing is a reliably efficient technique for the introduction of precise sequence changes in cultured cells via homologous recombination (9,10,11).

Cohesin complexes were purified via FLAG immunoprecipitation from nuclear extracts derived from parental HCT116 cells and each of the eleven cohesin endogenous epitope-tagged derivatives. Western blot was then performed with antibodies to FLAG and to each of the eleven known components of cohesin (Fig. S12). This experiment demonstrated that core subunits SMC1A, SMC3, STAG2, as well as PDS5A, are highly expressed at roughly equimolar levels; RAD21, STAG1, and WAPL are expressed at similar intermediate levels, and PDS5B, Sororin, NIPBL, and MAU2 are expressed at much lower levels. This experiment also demonstrated that SMC1A, SMC3, STAG2, and Sororin are most efficient at co-purifying the other known components of cohesin, and confirmed the known mutual exclusivity of STAG1 and STAG2 in cohesin complexes (12).

Dual affinity purification was then performed on nuclear extracts from the eleven epitope-tagged cell lines and parental HCT116 cells. Initially, affinity purifications were separated using SDS-PAGE and stained with silver, demonstrating that this purification approach made it possible to purify cohesin to high levels of homogeneity using each of the 11 known subunits as baits (Fig. S13). The protein composition of individual affinity purifications was then interrogated by GeLC-MS/MS following Coomassie staining, which results in sensitive protein identification by concentrating the sample and removing nonprotein contaminants (13). Proteins represented by two or more unique peptides in affinity purifications from epitope-tagged cells but absent in isogenic parental cells are listed in the eleven tabs of Table S2. All known components of cohesin (but no other known structural components of chromatin) were reciprocally identified in this analysis (Tables S2,S3), as well as other known cohesin interacting proteins such as components of the Mediator complex (14).

Protein-protein interaction networks were then identified using STRING (Fig. 1A) (15). This is, to our knowledge, the first entire protein complex/pathway whose interactome has been elucidated by endogenous tagging of each of its known components in human cells. The interaction network was by far the most highly enriched for proteins involved in RNA splicing, RNA binding, and/or which contained known RNA-recognition motifs than for any other pathway or protein domain (FDR 1.79 × 10^−42^) (Fig. 1B, Tables S4 and S5) (16). Particularly notable splicing factors included the proteins encoded by the SF3B1 oncogene, the RBM10 tumor suppressor gene, and the EFTUD2, SNRNP200, and PRPF31 components of the U4/U6.U5 tri-snRNP complex (17,18). These data were particularly intriguing in light of the recent observation that depletion of splicing factors in human cells can result in a paradoxical loss of sister chromatid cohesion (19,20,21,22).

**Fig. 1.**
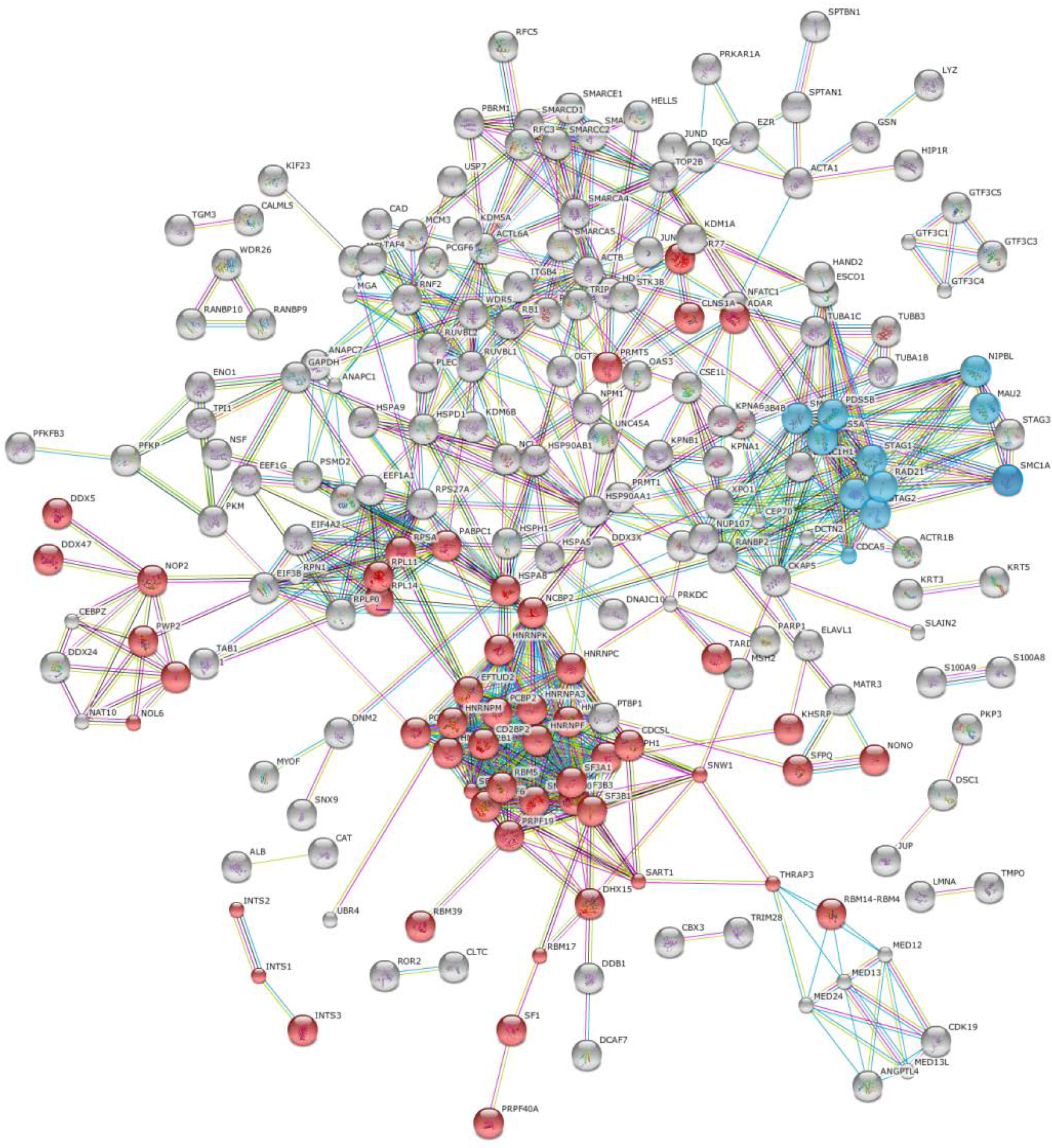
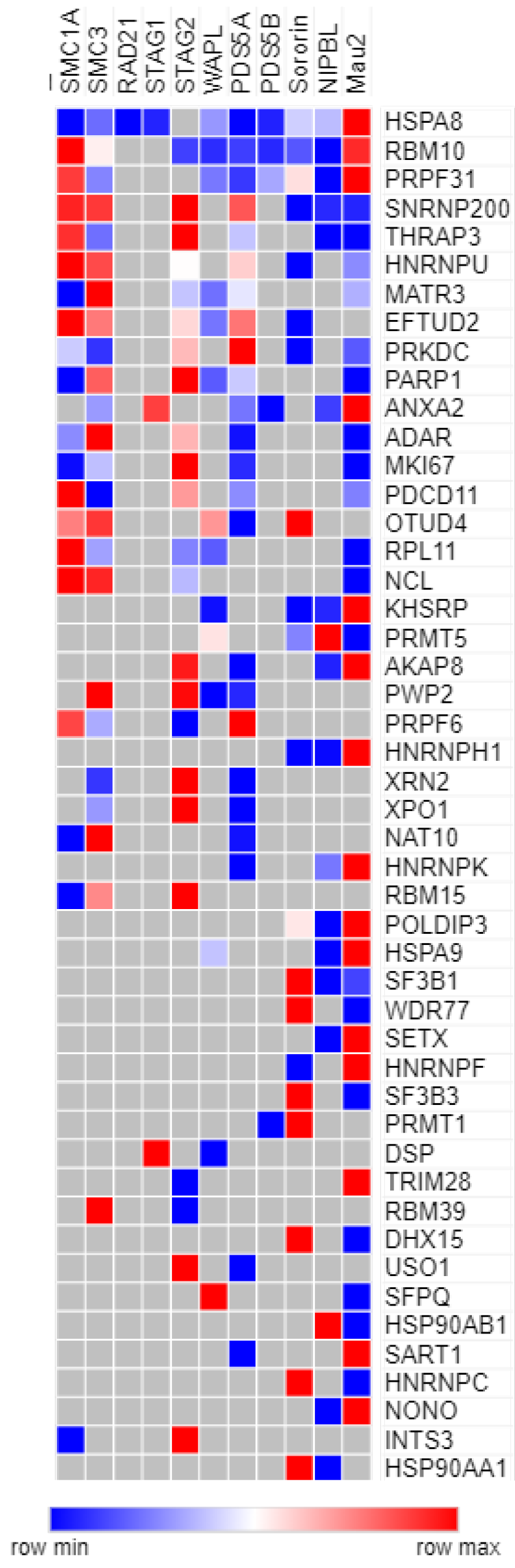
Cohesin interactome. **(A)** Mass spectrometry data was analyzed using STRING software for the identification of functional protein interaction networks. Nodes represent proteins identified by mass spectrometry from dual affinity purifications of endogenous epitope-tagged cells. Blue nodes represent each of the 11 cohesin subunits used as baits. Red nodes represent splicing factors and proteins with RNA binding domains as identified by GO, KEGG, and Pfam analysis. Edges represent protein-protein interactions, with teal and pink representing known interactions from curated databases and experimentally determined, respectively; green, red, and blue represent predicted interactions via gene neighborhoods, gene fusions, and gene co-occurrences, respectively; and light green, black, and violet represent interactions predicted by text mining, co-expression, and protein homology analysis, respectively. **(B)** Heatmap of interactions between cohesin subunits (columns) and interacting splicing factors/RNA binding proteins (rows). Proteins identified in two or more affinity purifications are shown. Colors as shown on the key represent the predicted relative abundance (ion area) of each protein in each of the eleven affinity purifications.

Other novel putative cohesin interactors identified in three or more dual affinity purifications included USP13 (deubiquitinase), MCM3 (replication licensing factor), MGA (myc-associated transcription factor), CKAP5 (kinetochore), and ANKRD11 (Cornelia de Lange Syndrome gene of unknown function). The details and functions of their interactions with cohesin will be described elsewhere.

Next, Western blot with antibodies to splicing factors and RNA binding proteins was performed on dual affinity purifications from nuclear extracts of parental HCT116 cells and SMC3 epitope-tagged derivatives (Fig. 2A). This experiment confirmed the interaction between endogenous cohesin and endogenous SF3B1, SF3B3, ADAR1, PRPF31, SNRNP200, EFTUD2, HNRNPU, RBM10, RBM15, HNRNPH, HSPA8, PDCD11, THRAP3, DDX47, and PRPF6. To confirm and generalize these findings, endogenous cohesin was immunoprecipitated with SMC1A antibodies from genetically unmodified HeLa cells and untransformed human epithelial cells (RPE-hTERT), and Western blot performed with a subset of these antibodies (Fig. 2B). Similar experiments were then performed on nuclear extracts that had been pre-treated with RNase and DNase, demonstrating that the interactions between cohesin and splicing factors/RNA binding proteins required neither a small nuclear RNA component nor chromatin (Fig. 2C).

**Fig. 2.**
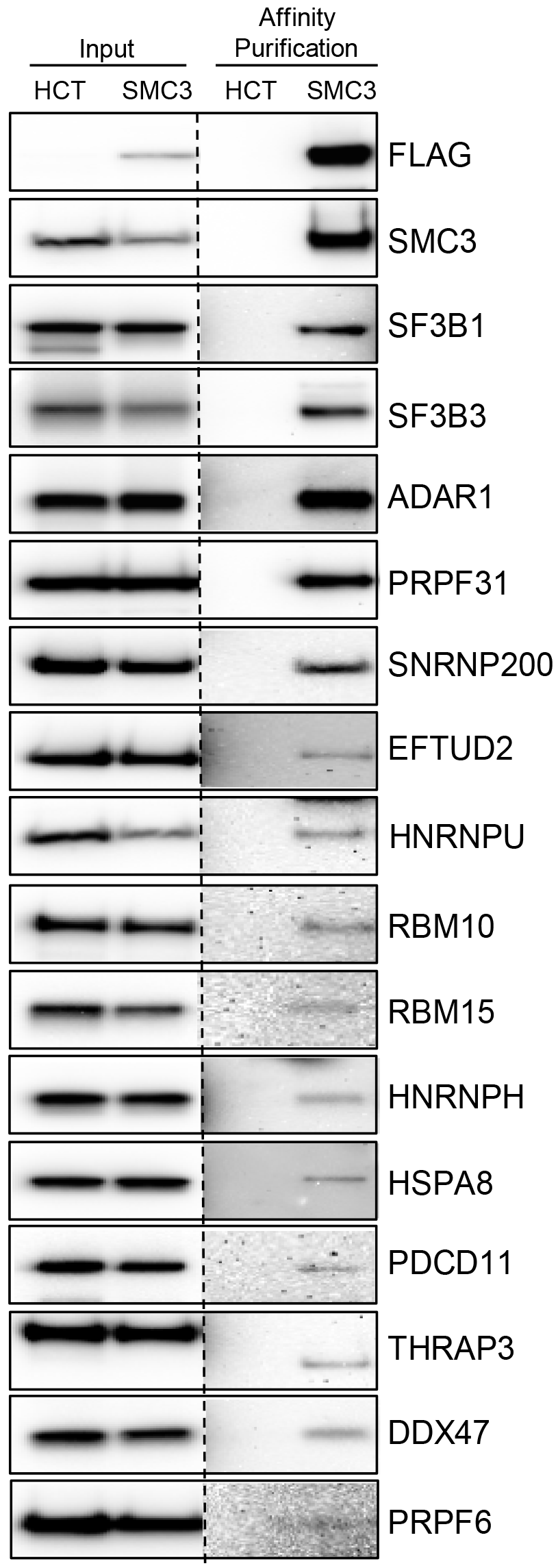
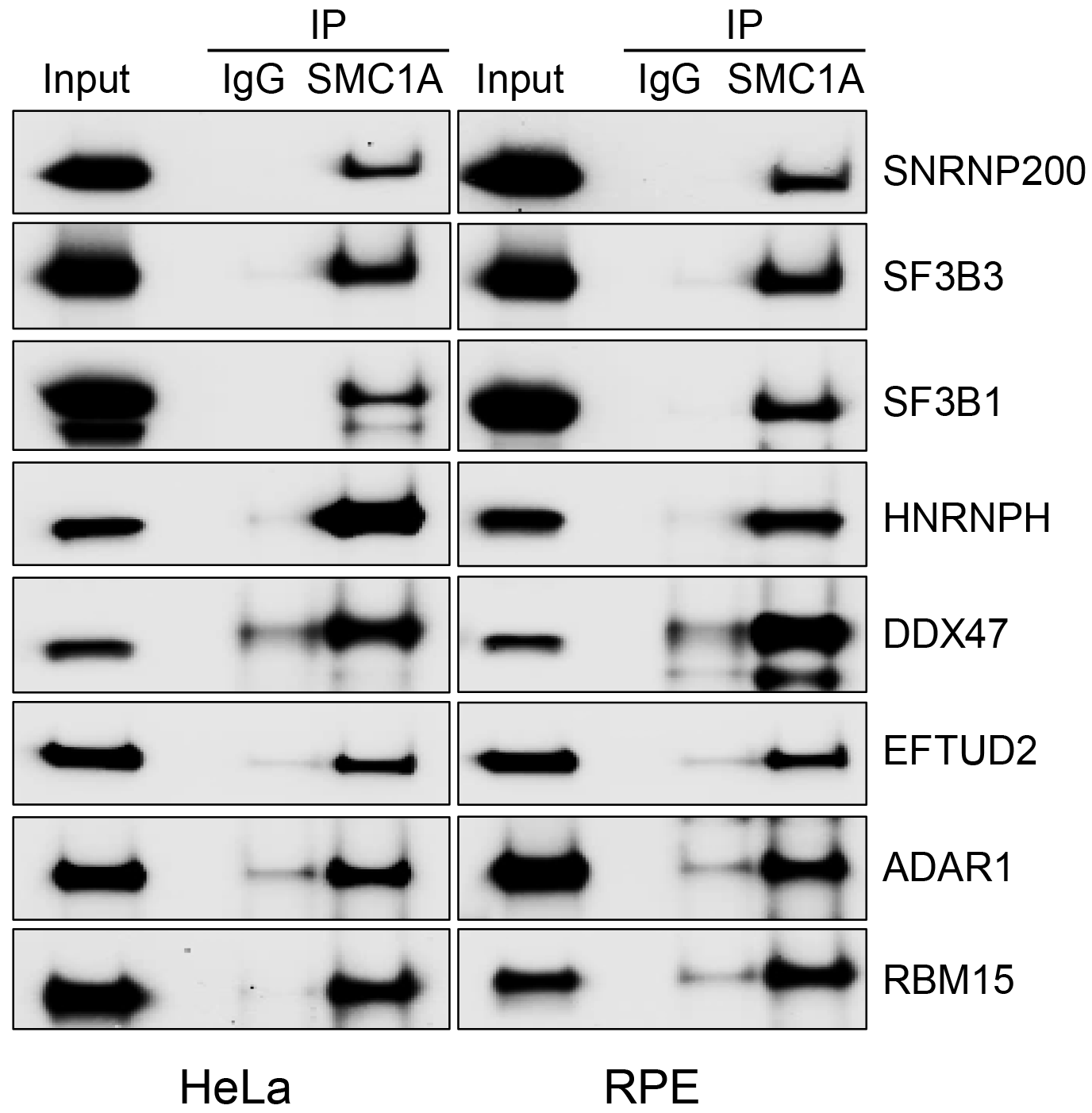
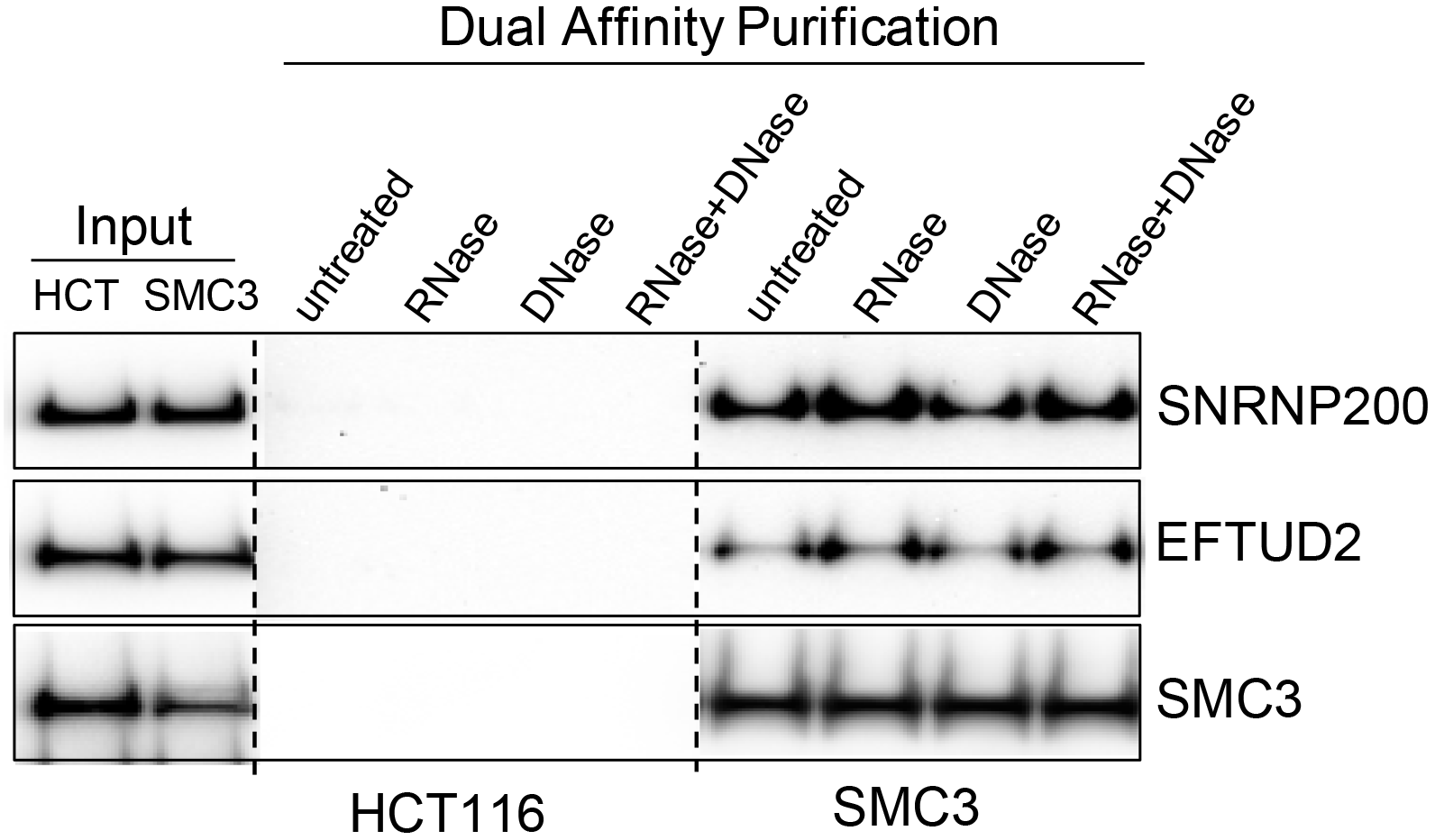
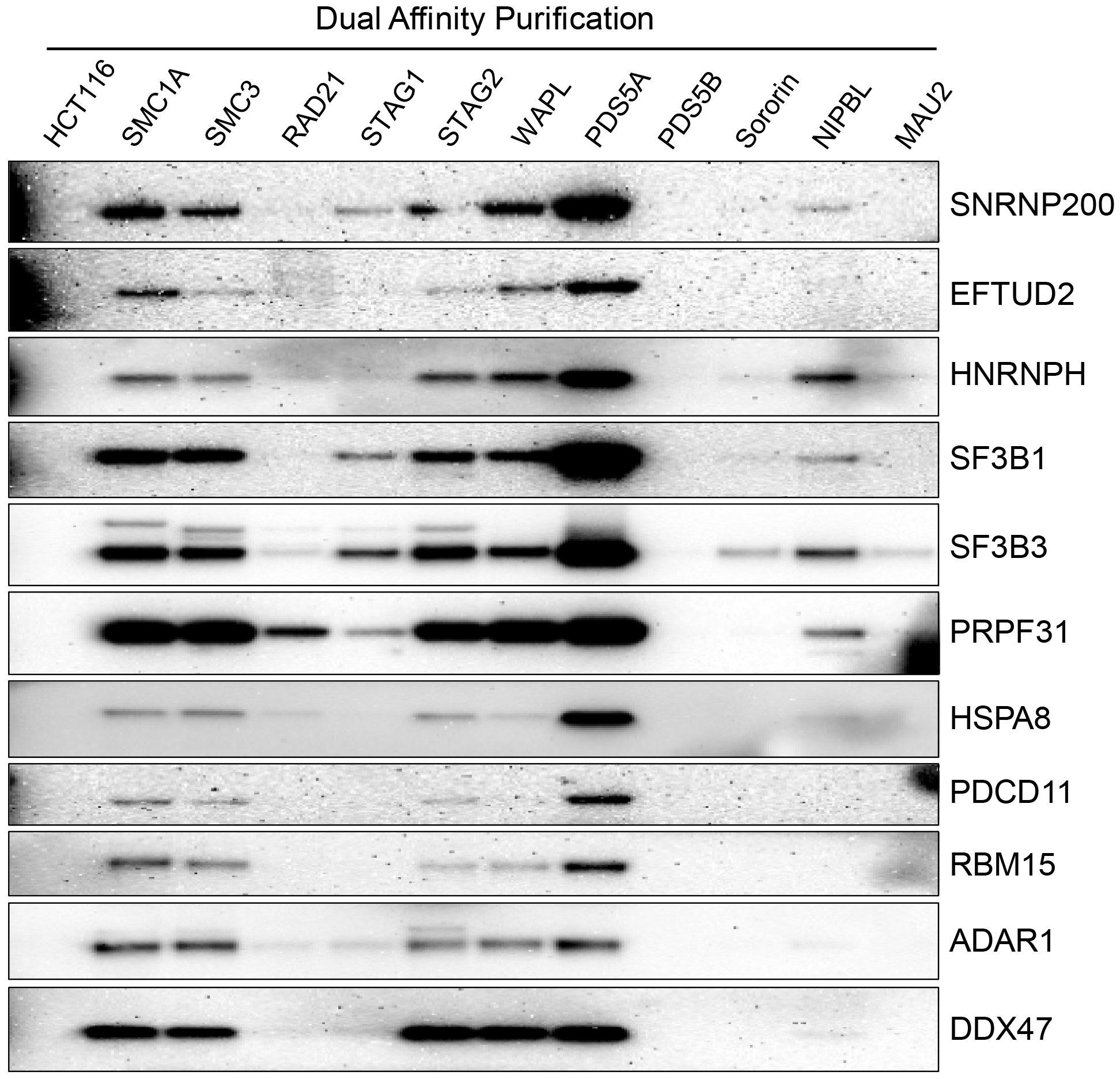
Interaction of cohesin with diverse splicing factors and RNA binding proteins. **(A)** Dual affinity purifications from nuclear extracts of HCT116 cells and SMC3 epitope-tagged derivatives were separated by SDS-PAGE and Western blot performed with the antibodies indicated. **(B)** Immunoprecipitations with IgG and SMC1A antibodies from HeLa and RPE-hTERT nuclear extracts were separated by SDS-PAGE and Western blot performed with the antibodies indicated. **(C)** Dual affinity purifications from RNase and DNase-treated nuclear extracts of HCT116 cells and SMC3 epitope-tagged derivatives were separated by SDS-PAGE and Western blot performed with the antibodies indicated. **(D)** Dual affinity purifications from nuclear extracts of HCT116 cells and eleven cohesin epitope-tagged derivatives were separated by SDS-PAGE and Western blot performed with the antibodies indicated.

To determine if splicing factors interact preferentially with cohesin complexes defined by the presence of specific cohesin subunits, Western blot with splicing factor antibodies was performed on dual affinity purifications from parental HCT116 cells and each of the 11 epitope-tagged cell lines (Fig. 2D). This experiment demonstrated that splicing factors/RNA binding proteins were most efficiently co-purified with PDS5A-containing cohesin complexes.

Since it has been shown that many splicing factors are paradoxically required for cell cycle progression (23,24,25), we next tested whether the interactions between cohesin and splicing factors were cell cycle regulated. To do this, HeLa cells were synchronized by double thymidine block, released, and whole cell lysates prepared from cells at different stages of the cell cycle. Next, cohesin was immunoprecipitated with SMC1A antibodies and splicing factor Western blot performed, demonstrating that the interaction between cohesin and splicing factors is substantially enhanced during mitosis (Fig. 3A,B, Fig. S14). Identical results were obtained with cells that had been arrested by hydroxyurea, RO-3306 (26), and nocodazole in the S, G2, and M phases of the cell cycle, respectively (Fig. S15).

**Fig. 3.**
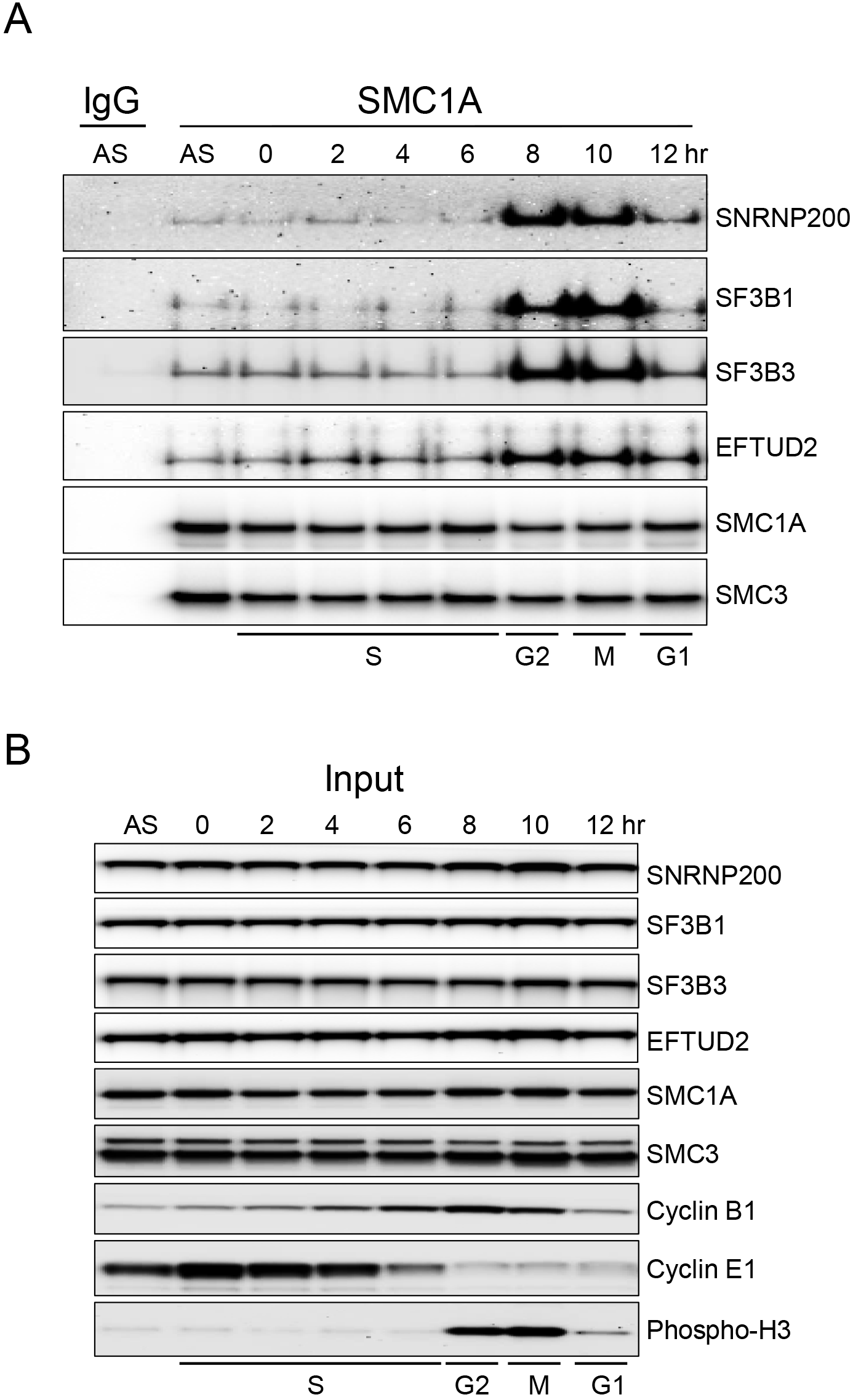
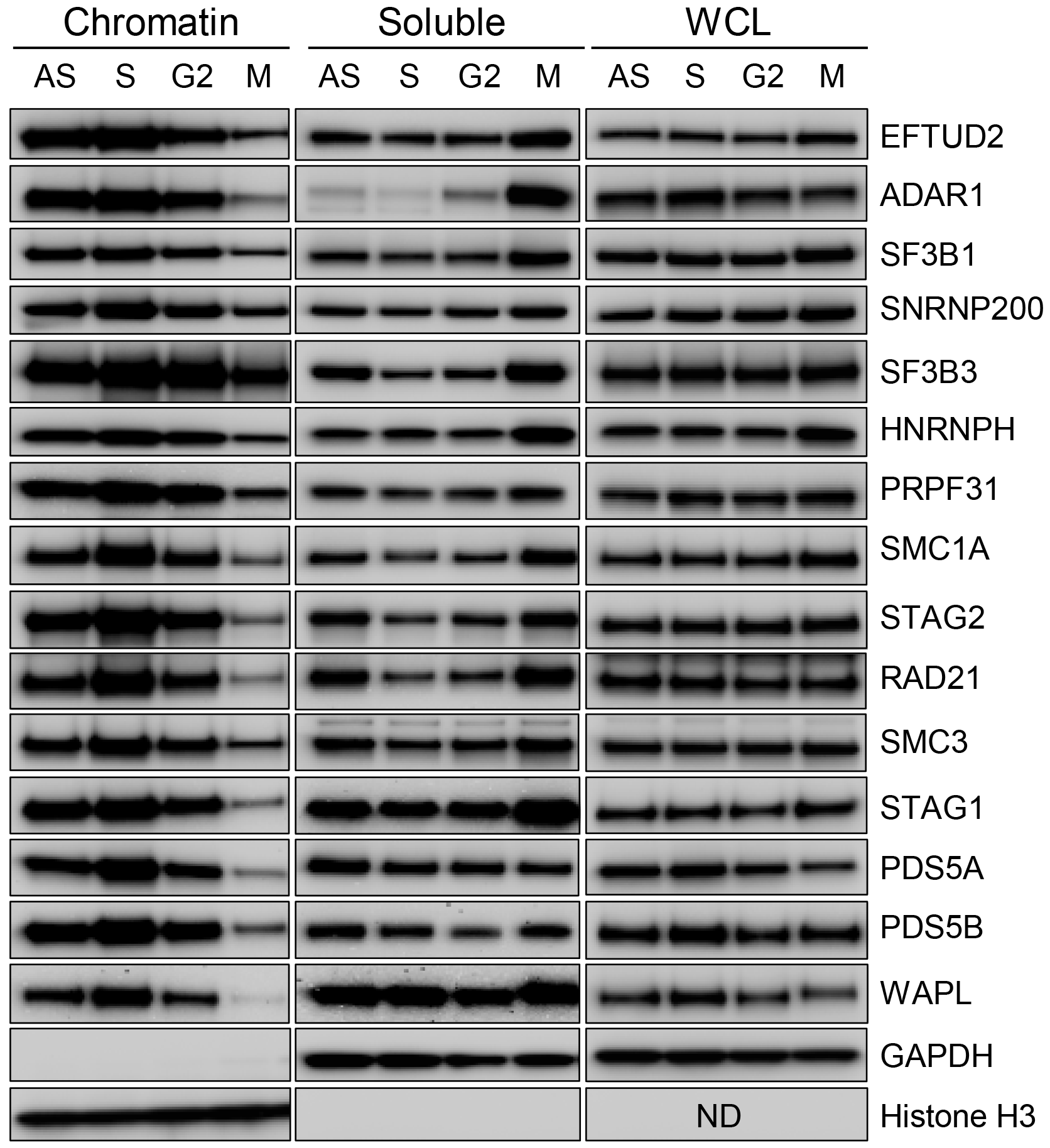
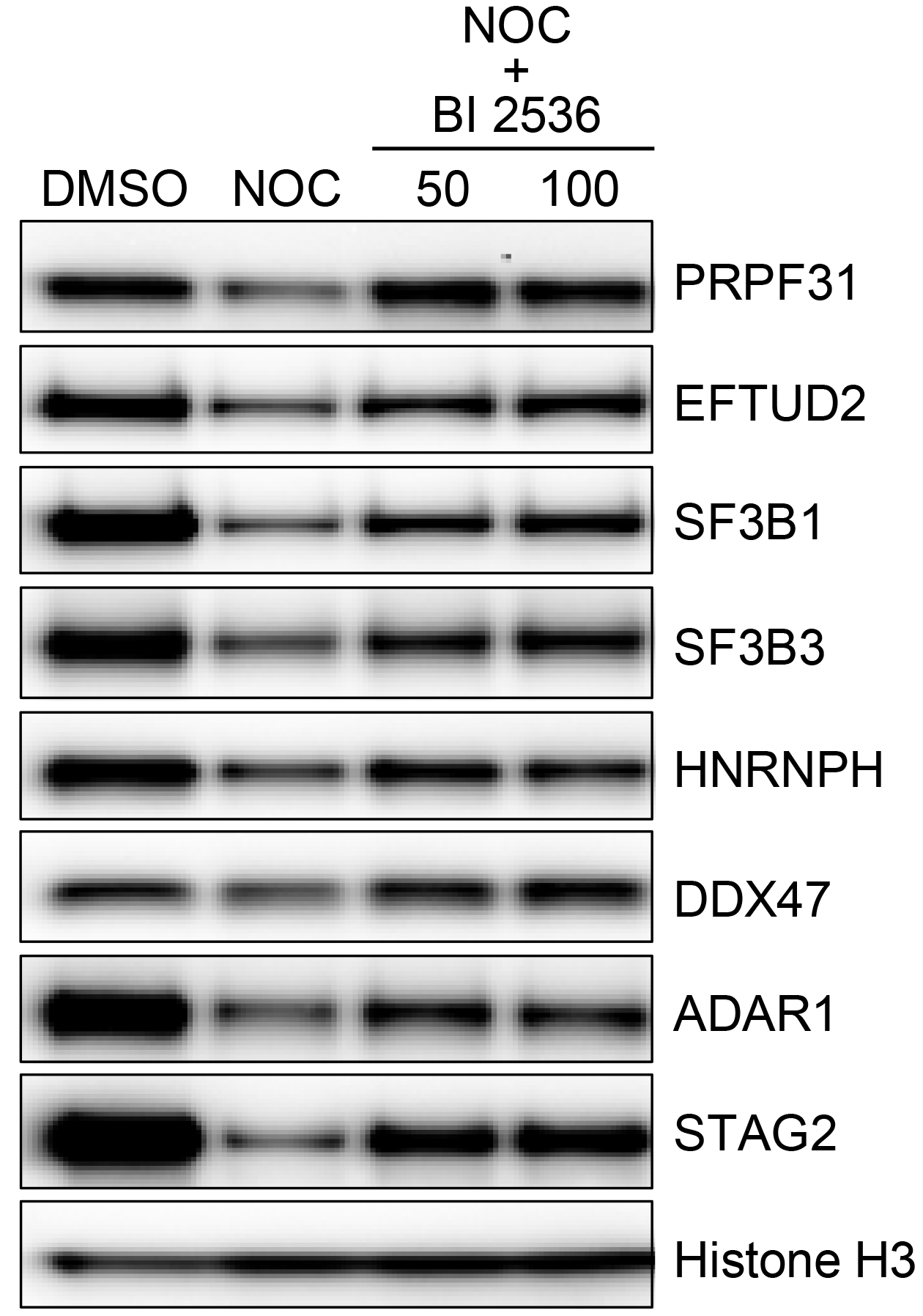
Cell cycle regulation of the interaction of splicing factors with cohesin and chromatin. **(A)** HeLa cells were synchronized by double thymidine block and whole cell protein lysates (1% NP40) prepared at different time points following release. Endogenous cohesin complexes were then immunoprecipitated with SMC1A antibodies or IgG control antibodies, and Western blot with the antibodies indicated was performed. **(B)** Direct Western blot of the lysates studied in (A), demonstrating that the amount of cohesin and splicing factor proteins in the lysates does not change over the course of the cell cycle, and demonstrating the efficiency of synchronization using cell cycle-stage specific antibodies (S phase – cyclin E1, G2 - cyclin B1, M – phospho-H3). **(C)** Proliferating HeLa cells were treated with DMSO (AS), hydroxyurea (S), RO-3306 (G2), and nocodazole (M) for 24 hours to arrest cells at different stages of the cell cycle. Chromatin, soluble, and whole cell lysates (WCL;RIPA) were prepared and studied by Western blot with the antibodies indicated. **(D)** HeLa cells were treated with 100 ng/mL nocodazole (NOC) for 24 hr., then treated with the PLK1 inhibitor BI 2536 (31) at 50 and 100 nM for 3 hr. Cells were then harvested, chromatin lysates prepared, and Western blot performed with the antibodies indicated.

Cohesin interacts robustly with chromatin during DNA replication and is released from chromatin during mitosis in a process known as the “cohesin cycle” (27). To test whether the interaction of splicing factors/RNA binding proteins with chromatin was similarly cell cycle regulated, HeLa cells were treated with hydroxurea, RO-3306, and nocodazole to arrest cells in S phase, G2, and mitosis, respectively, and chromatin, soluble, and total protein lysates were prepared. As expected, Western blot confirmed that cohesin was enriched on chromatin during DNA replication and released from chromatin during mitosis (Fig. 3C). Remarkably, splicing factors EFTUD2, ADAR1, SF3B1, SNRNP200, SF3B3, HNRNPH, and PRPF31 demonstrated a similar cell cycle-regulated interaction with chromatin. To generalize these findings, similar experiments were performed in RPE cells and HCT116 cells, with identical results (Fig. S16,S17).

In mitosis, the majority of cohesin is released from chromatin in prophase in a PLK1-dependent fashion known as the “prophase pathway” (28,29,30). To determine if the release of splicing factors from chromatin in mitosis is similarly PLK1-dependent, nocodazole-treated HeLa cells were incubated in the presence or absence of PLK1 inhibitors (31) and chromatin extracts were prepared. Western blot with antibodies to cohesin subunits and splicing factors demonstrated that the release of splicing factors from chromatin during mitosis and the release of cohesin from chromatin during mitosis were equivalently PLK1-dependent (Fig. 3D). Together these data show that the interaction between diverse splicing factors and chromatin is cell cycle regulated, and that the interaction of splicing factors with chromatin follows the cohesin cycle and the prophase pathway. This finding supports and extends recent studies emphasizing the important functional roles of the interaction of splicing factors with chromatin (32,33).

To determine if depletion of specific cohesin-interacting splicing factors results in cell cycle phenotypes, HeLa H2B-GFP cells were transfected with multiple independent, validated siRNAs (Fig. S18) for seven different cohesin-interacting splicing factors/RNA binding proteins and imaged every five minutes for 36 hours using a Leica SP8 laser scanning confocal microscope (Movie S1,S2). Depletion of EFTUD2, SNRNP200, and PRPF31, three different cohesin-interacting components of the U4/U6.U5 tri-snRNP complex, as well as SF3B3, resulted in a stereotyped prometaphase arrest characterized by an inability of sister chromatids to maintain alignment at the metaphase plate, and a failure to successfully execute anaphase (Fig. 4A,B). This arrest is phenotypically similar to the mitotic arrest caused by depletion of Sororin (Fig. 4B). Depletion of SF3B1 resulted in an immediate, completely penetrant interphase cell cycle arrest (Movie S2).

**Fig. 4.**
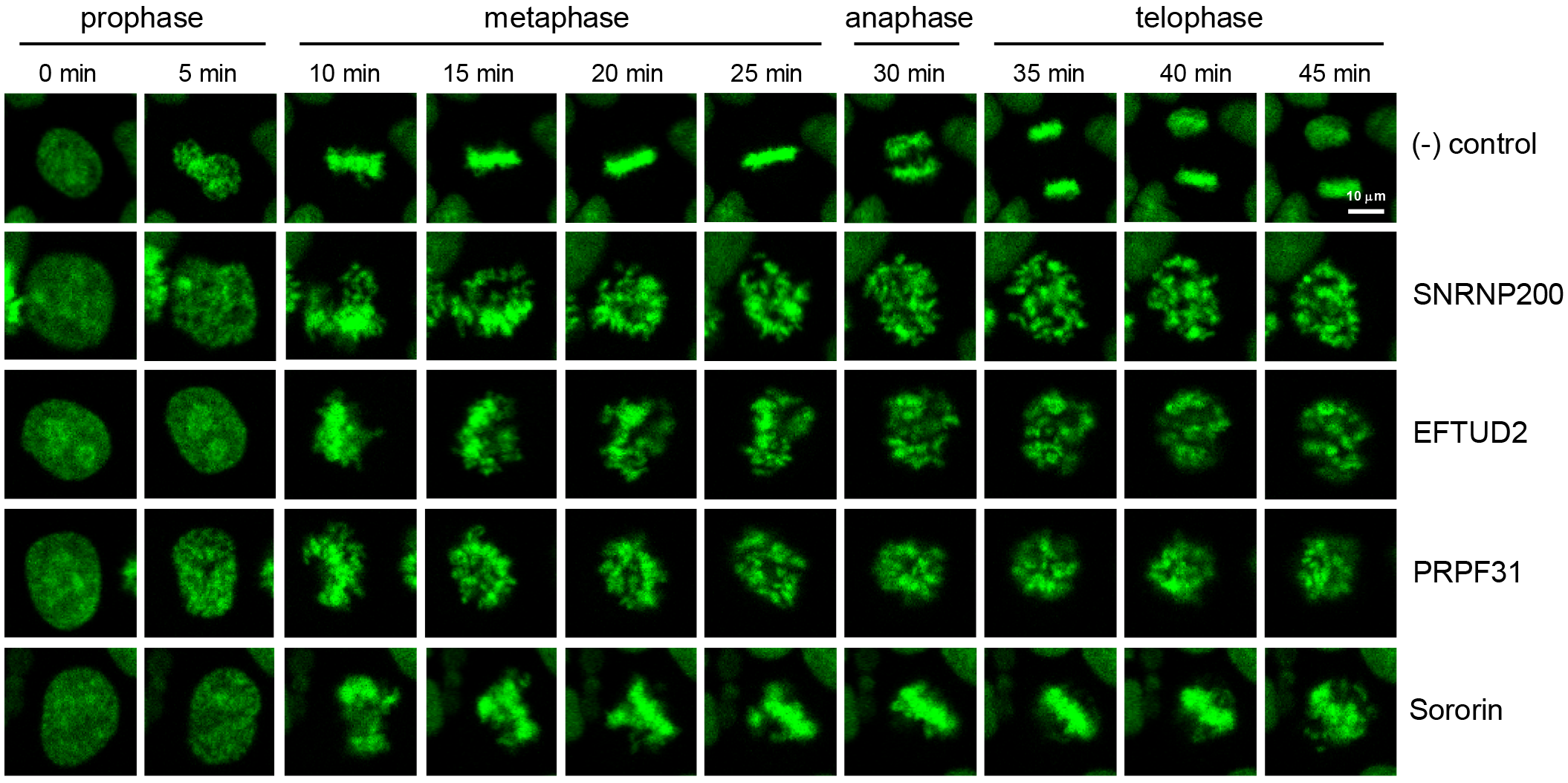
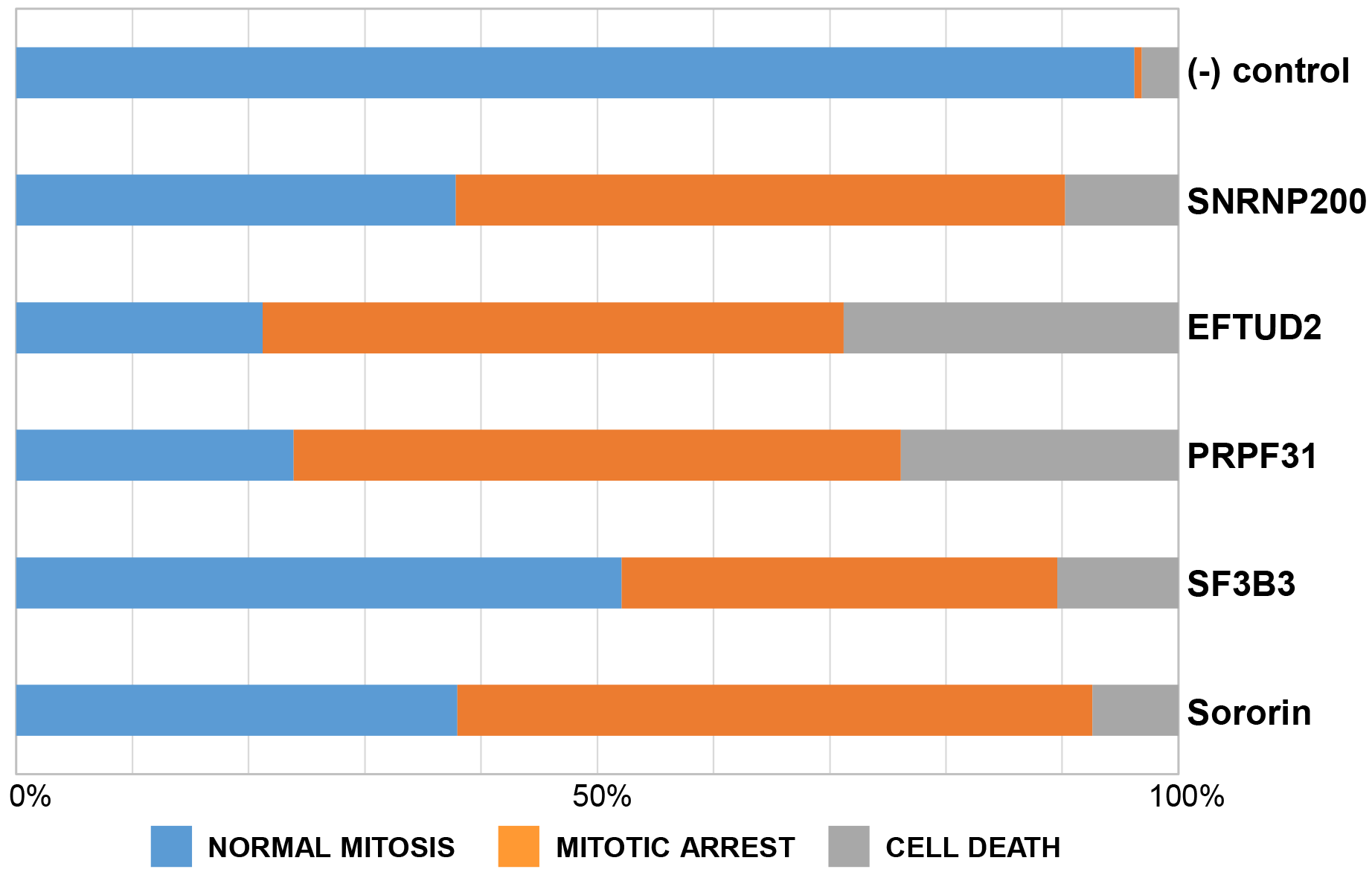
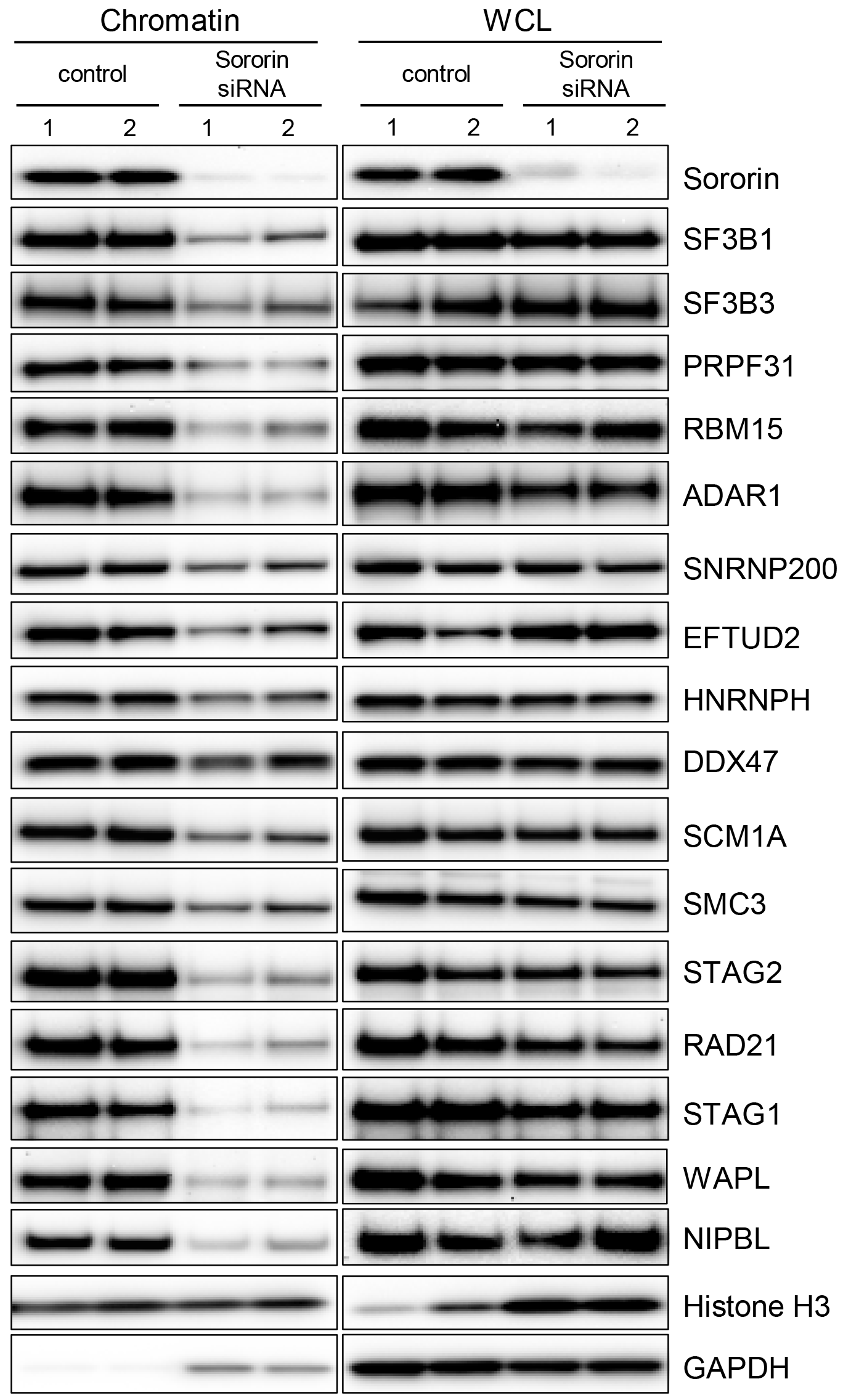
Cohesin-interacting splicing factors are required for cell cycle progression. HeLa-H2B-GFP cells were transfected with siRNAs to cohesin-interacting splicing factors and studied by live cell imaging as described in Materials and Methods. Representative aberrant mitoses in siRNA-transfected cells are shown in **(A)**. For complete movies with images every 5 min. from 36-72 hr. post- transfection, see Supplemental Movies S1 and S2. **(B)** Quantification of aberrant mitoses in siRNA-transfected cells. Cells in five fields were counted, representing 48-158 cells, depending on the transfected siRNA. **(C)** HeLa cells were transfected with control (#1-untransfected, #2-control siRNA) and sororin siRNAs (#1-s535459, #2-s535461). Three days after transfection, chromatin lysates and whole cell lysates (WCL; RIPA) were prepared and Western blot performed with the antibodies indicated.

Next, we tested whether these cell cycle arrests were accompanied by alterations in the interaction of cohesin with chromatin. To do this, chromatin and whole cell lysates were prepared from siRNA-transfected cells, and Western blot with antibodies to cohesin and splicing factors was performed. Depletion of Sororin was accompanied by a marked decrease in the levels of chromatin bound splicing factors (Fig. 4C). Depletion of SF3B1 was accompanied by a decrease in the levels of chromatin-bound and total cohesin, except for a marked increase in Sororin (Fig. S19). Depletion of SNRNP200 resulted in a decrease in the levels of chromatin-bound and total STAG1 and NIPBL (Fig. S20). Together these data support the hypothesis that levels of splicing factors and cohesin are co-regulated and depend on each other for their interaction with chromatin.

Finally, we hypothesized that cohesin-interacting splicing factors, like cohesin itself (34,35,36,37), could regulate genomic organization. To provide proof-of-principle, DNase Hi-C (38) was performed on biological replicates of HeLa cells transfected with either scrambled siRNA or SF3B1 siRNA. Bioinformatics analysis identified two specific roles for SF3B1 in the maintenance of chromosome compartment structure. First, depletion of SF3B1 resulted in the “weakening” of compartment structure, in that the segregation between active “A” compartments and inactive “B” compartments was less pronounced in SF3B1-depleted cells than in control cells. This weakening of compartment structure can be visualized as a decrease in the magnitude of the observed Hi-C correlation values in control and SF3B1-depleted cells, depicted in Figs. 5A, 5B, and S21 and described in detail in the legend. Second, in addition to the weakening of compartment structure, depletion of SF3B1 resulted in the transformation of B compartments into A compartments, an effect that was particularly pronounced in larger chromosomes. This transformation of B compartments into A compartments is depicted in the eigenvector plots shown in the bottom of Fig. 5A and Fig. S21 and described in detail in the legend. As B compartments are transformed into A compartments, the total number of compartments in ten of the largest chromosomes decreases (Fig. 5C), and the total length of A compartments increases (Fig. 5D). In contrast to this alteration in compartment structure, topologically associated domains (TADs) are preserved in SF3B1-depleted cells. This maintenance of TAD structure is depicted in Fig. 5E, 5F, and S22 and described in detail in the legend. This experiment provides the first evidence pointing to a role for a splicing factor in the maintenance of genomic organization.

**Fig. 5.**
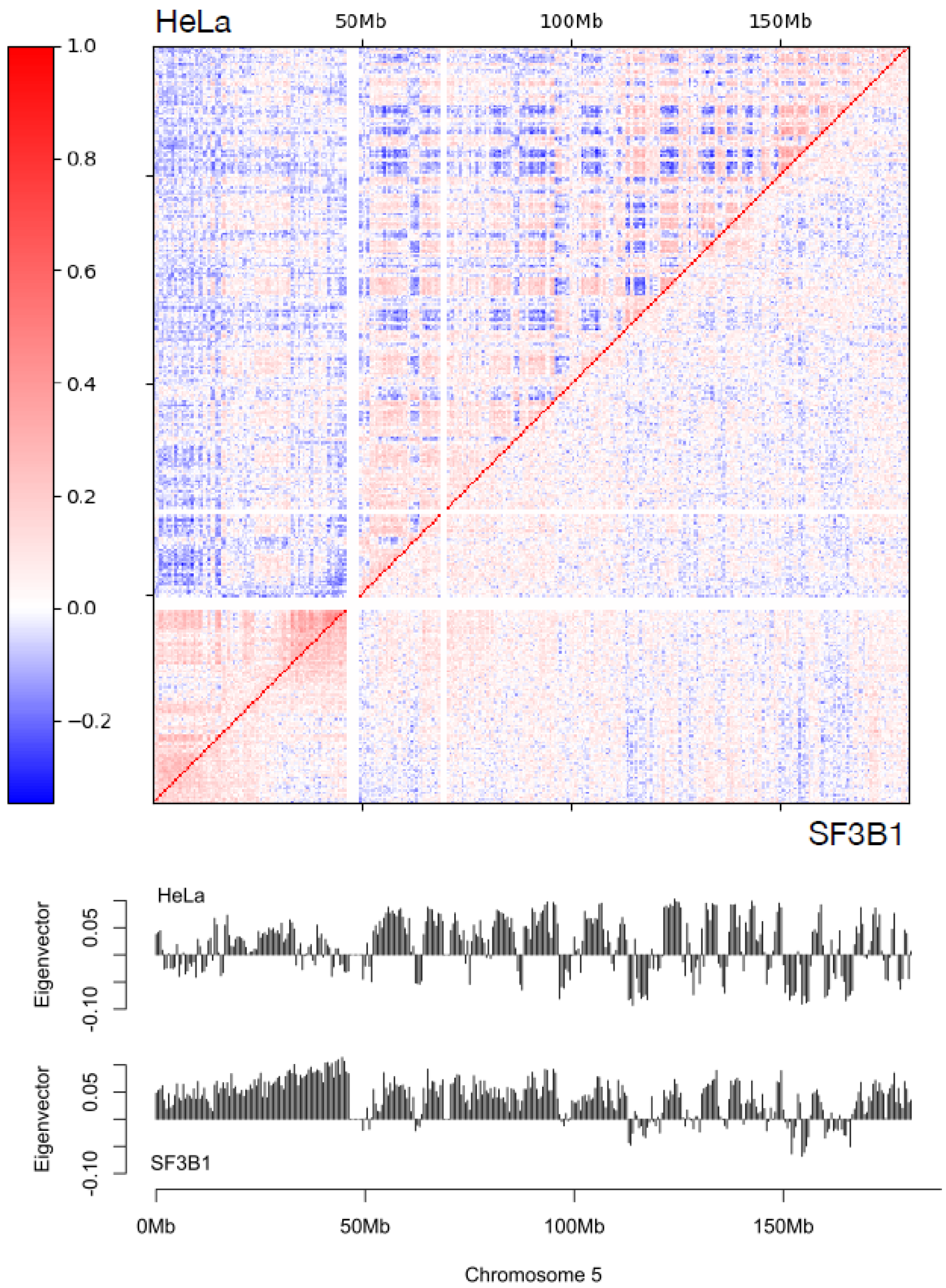
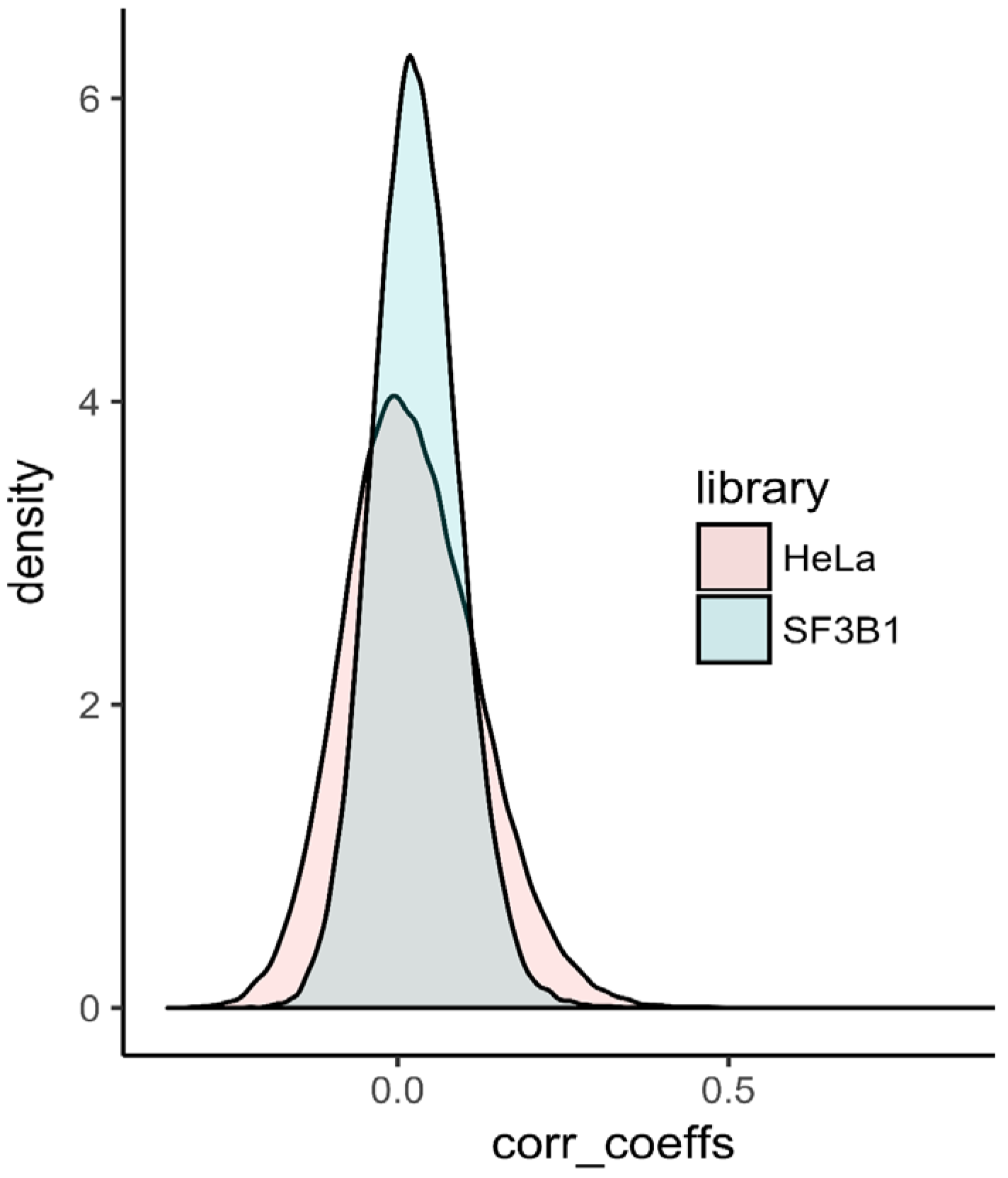
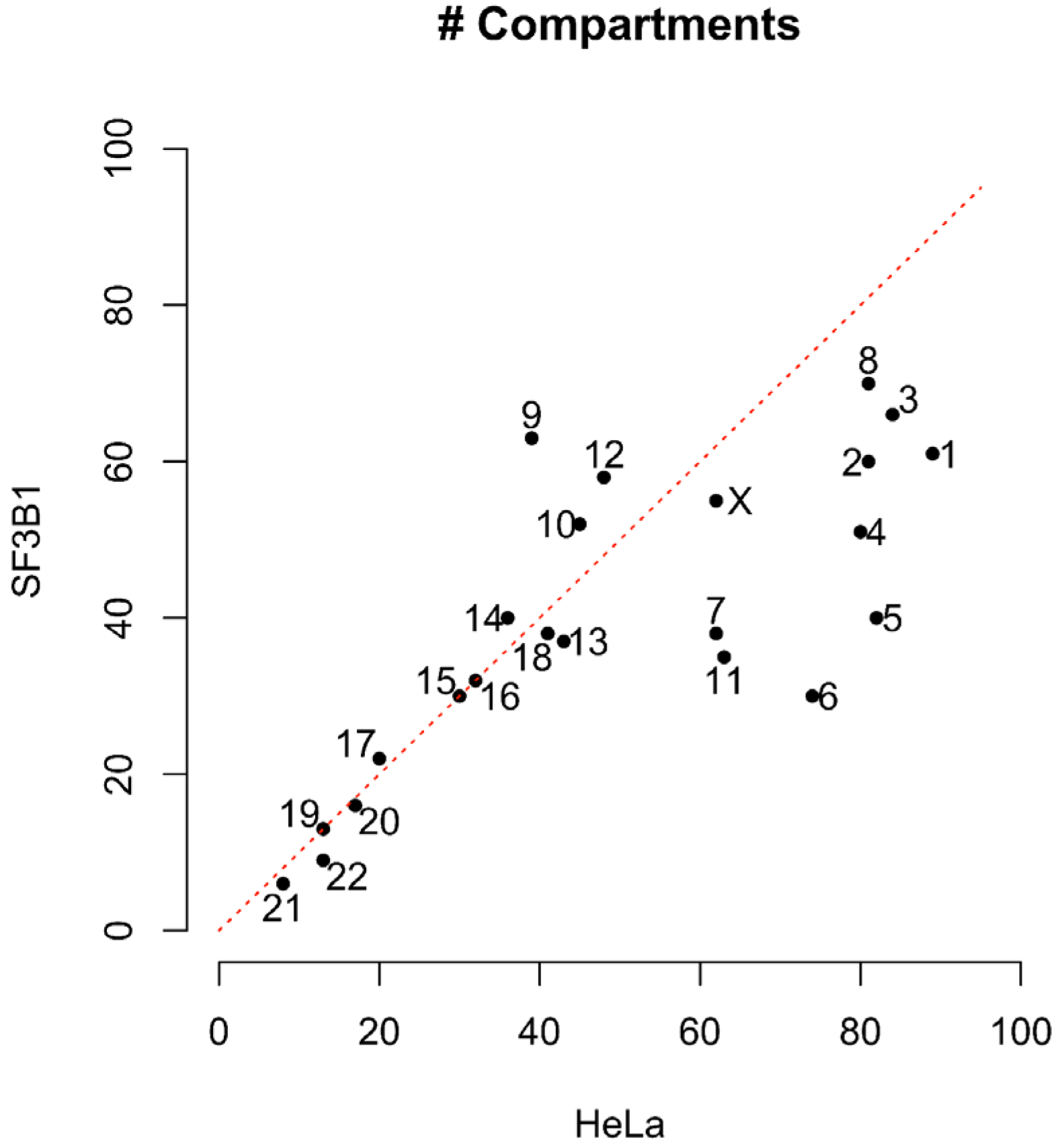
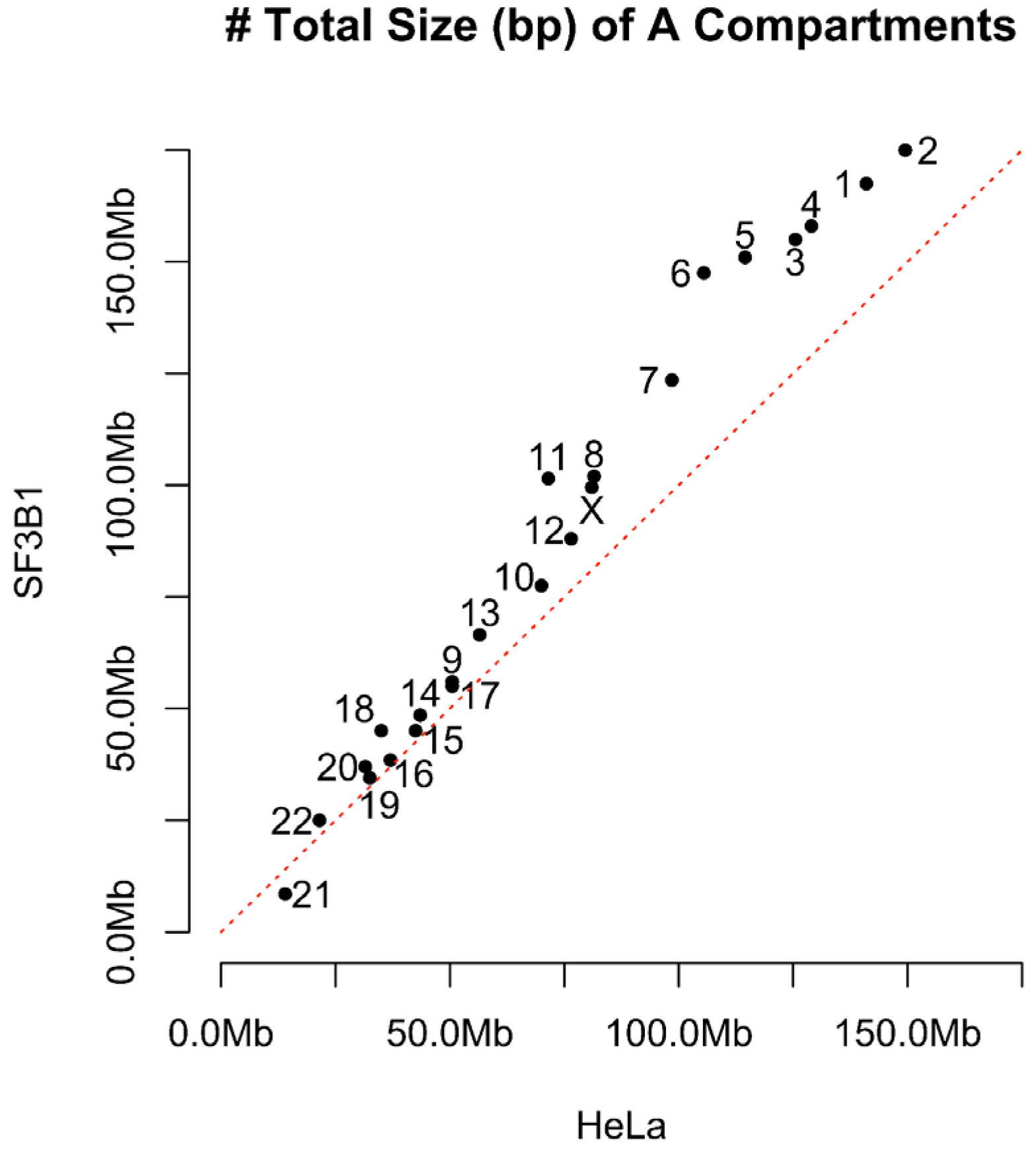
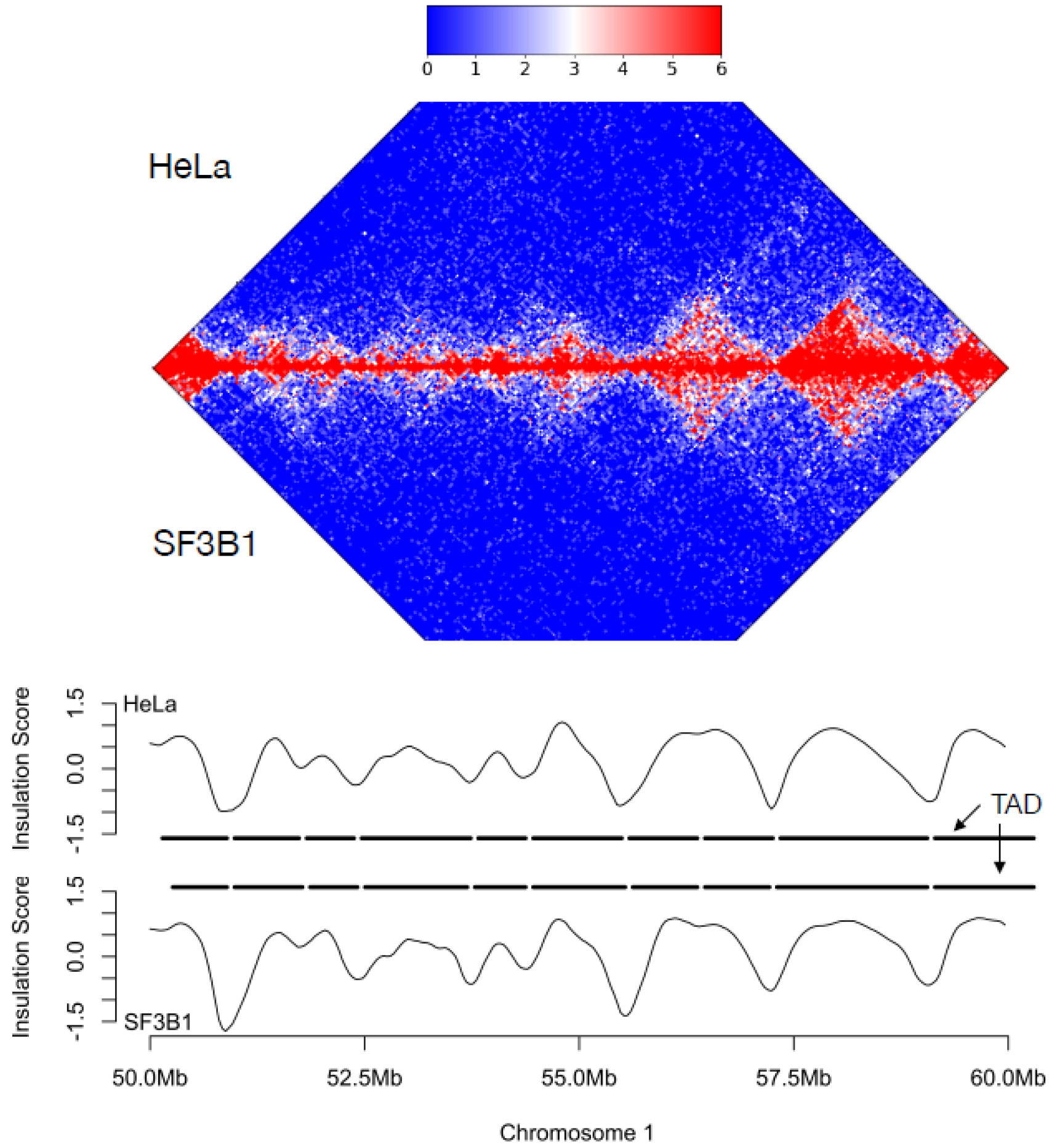
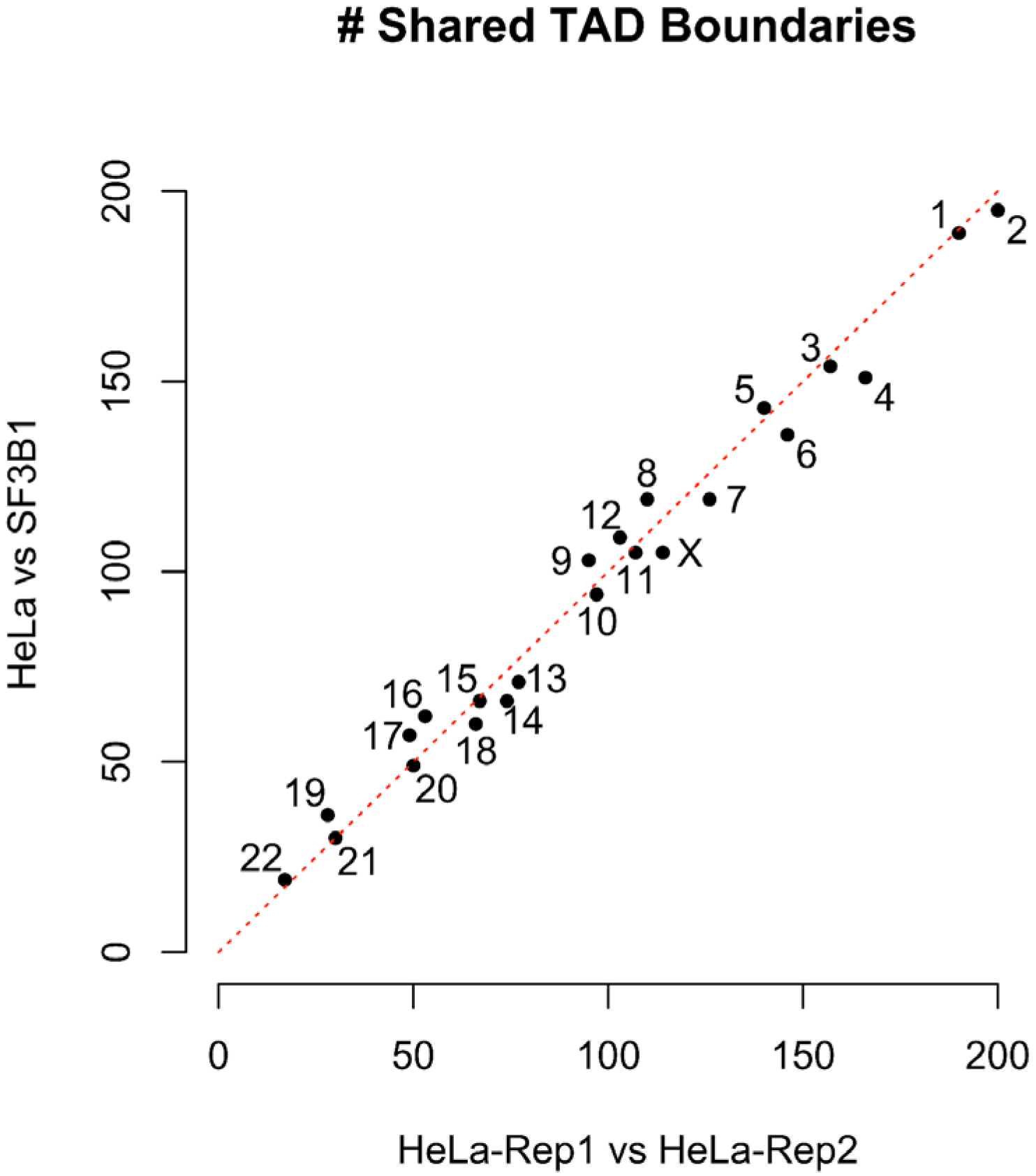
Consequences of SF3B1 depletion on 3D genomic organization. Panels A-D demonstrate alterations of compartment structure in SF3B1-depleted cells, and panels E, F demonstrate retention of topologically associated domains (TADs) in SF3B1-depleted cells. **(A)** Top: Loss of compartment structure is seen in the Hi-C correlation matrix for chromosome 5, with compartments visually apparent as a checkerboard pattern in HeLa cells transfected with negative control scrambled siRNA (upper left triangle) and reduced in HeLa cells transfected with SF3B1 siRNA (lower right triangle). Bottom: Compartmentalization is quantified by eigenvectors derived from the two Hi-C matrices, with fewer distinct compartments in HeLa cells transfected with SF3B1 siRNA than in cells transfected with scrambled siRNA. The active A compartment corresponds to eigenvector values >0. **(B)** The decrease in extreme (both high and low) correlation values, which is visually apparent in the heat maps in panel (A), is visualized directly here. The panel plots overlapping histograms of the Hi-C correlation values for chromosome 5 in HeLa cells transfected with SF3B1 siRNA (pink) versus cells transfected with scrambled siRNA (blue), showing that the latter exhibits more extreme correlations, corresponding to more pronounced compartment structure. **(C)** The number of compartments per chromosome is plotted for HeLa transfected with scrambled siRNA (x-axis) versus HeLa cells transfected with SF3B1 siRNA (y-axis). The number of compartments is reduced in SF3B1-depleted cells in the largest chromosomes. **(D)** The total size of A compartments per chromosome is plotted for HeLa transfected with scrambled siRNA (x-axis) versus HeLa cells transfected with SF3B1 siRNA (y-axis). The total size of A compartments is increased in SF3B1-depleted cells in the largest chromosomes, demonstrating that the number of compartments is reduced because B compartments have become A compartments. **(E)** TADs are consistent between control transfected HeLa cells and SF3B1 siRNA transfected HeLa cells. Top: Normalized Hi-C contacts for a 10 Mbp region of chromosome 1 are shown, focusing on intrachromosomal contacts. Contacts for control transfected HeLa appear above the x-axis, mirrored with corresponding contacts in cells transfected with SF3B1 siRNA. Consequently, TADs appear as red diamonds of enriched local contacts. Bottom: Insulation scores and TADs, called on the same region from HeLa cells and SF3B1 depleted HeLa cells, are highly concordant. **(F)** The figure plots, for each chromosome, the number of TAD boundaries shared between the two replicates of HeLa (x-axis) versus HeLa and SF3B1 (y-axis), demonstrating that TAD boundaries are unchanged after depletion of SF3B1.

Here we show that in addition to their well-known canonical role in cell biology, diverse splicing factors are among the most ubiquitous cohesin interacting proteins, interact with cohesin and chromatin in a cell cycle regulated fashion, and like cohesin can be required for maintenance of genomic organization. These findings identify a new non-canonical function for splicing factors and RNA binding proteins, provide mechanistic insight into the mysterious role of splicing factors and RNA binding proteins in sister chromatid cohesion and cell cycle progression, and have potential implications in explaining the mechanistic role of splicing factor mutations in diverse human cancers and inherited disease states.

## Acknowledgements

Mass spectrometry and bioinformatics analysis was conducted at the Mass Spectrometry and Proteomics Resource Laboratory, FAS Division of Science, Harvard University. DNA sequencing for Hi-C was performed at the Institute for Genome Sciences, University of Maryland School of Medicine. This research was supported by the National Institutes of Health (grants R01CA169345 and R21CA143282 to T.W.), Alex’s Lemonade Stand (T.W.), the Hyundai Hope on Wheels Foundation (T.W.), and the NIH Common Fund 4D Nucleome Project (grant U54DK107979 to W.S.N.). The Lombardi Comprehensive Cancer Center is supported by NIH grant P30CA51008. We thank Luke Tallon and Lisa Sadzewicz for assistance for DNA sequencing, Peter Johnson for assistance with live cell imaging, Ahmad Daher and Frederick Ghandchi for assistance with gene editing, and David Sabatini for critical comments on the manuscript.

## Supplementary Materials

Materials and Methods

Figs. S1 to S22

Tables S1-S6

Movies S1, S2

